## Supplementary Figures for "Structural screens identify candidate human homologs of insect chemoreceptors and cryptic *Drosophila* gustatory receptor-like proteins"

Figure 2—figure supplement 1

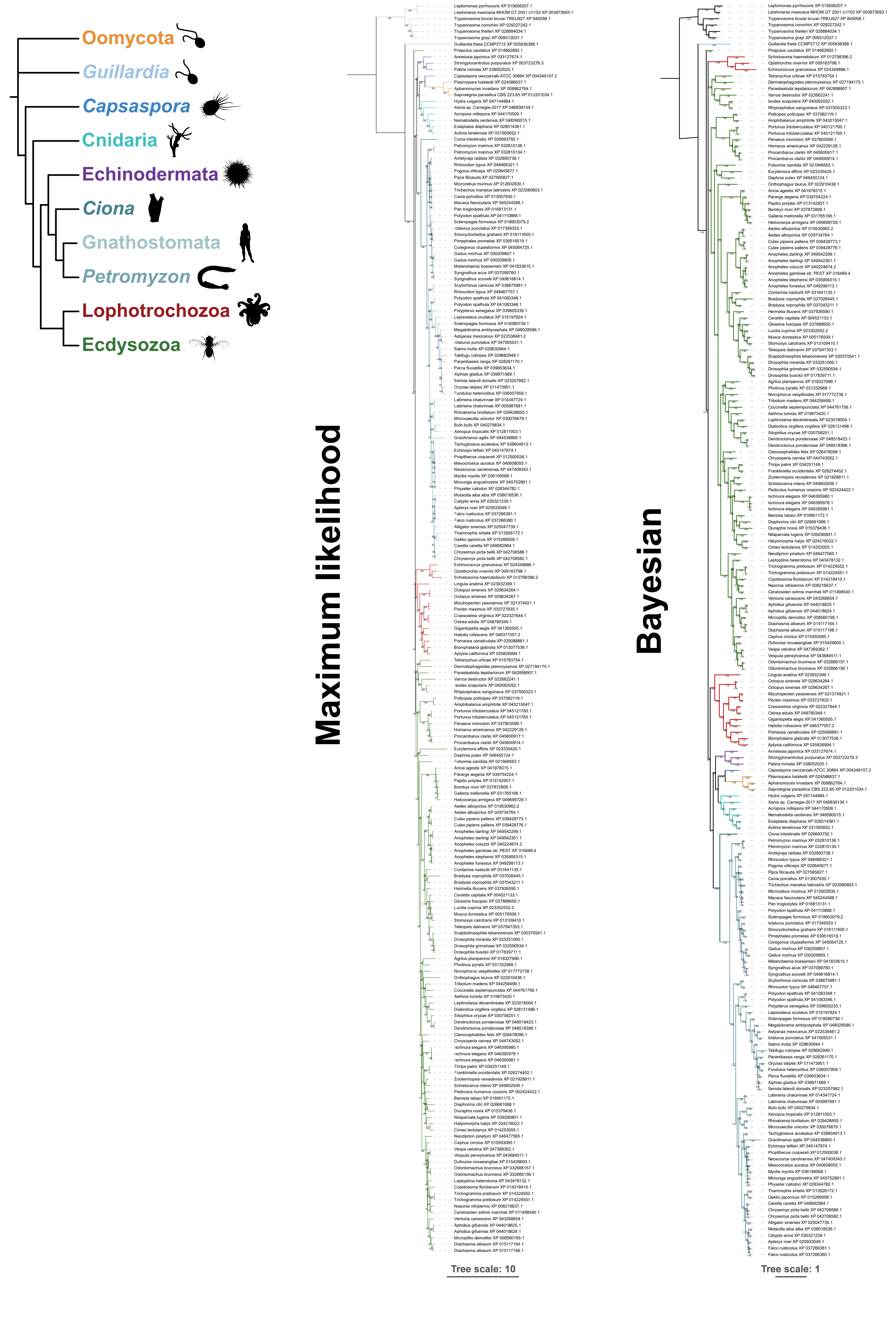

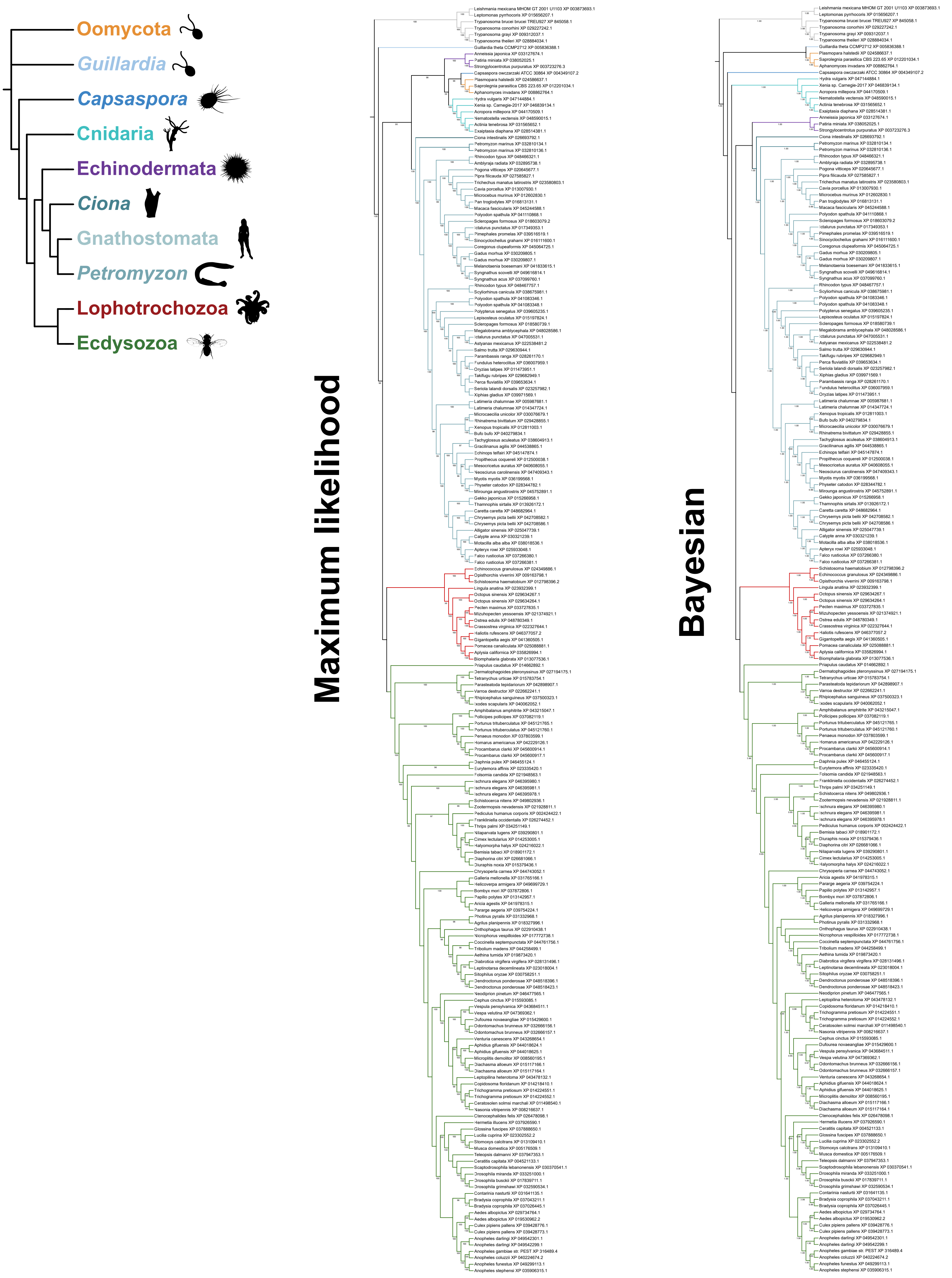
Oomycota
Guillardia
Capsaspora
Cnidaria
Echinodermata
Ciona
Gnathostomata
Petromyzon
Lophotrochozoa
Ecdysozoa
Maximum likelihood
Bayesian

Figure 2—figure supplement 3

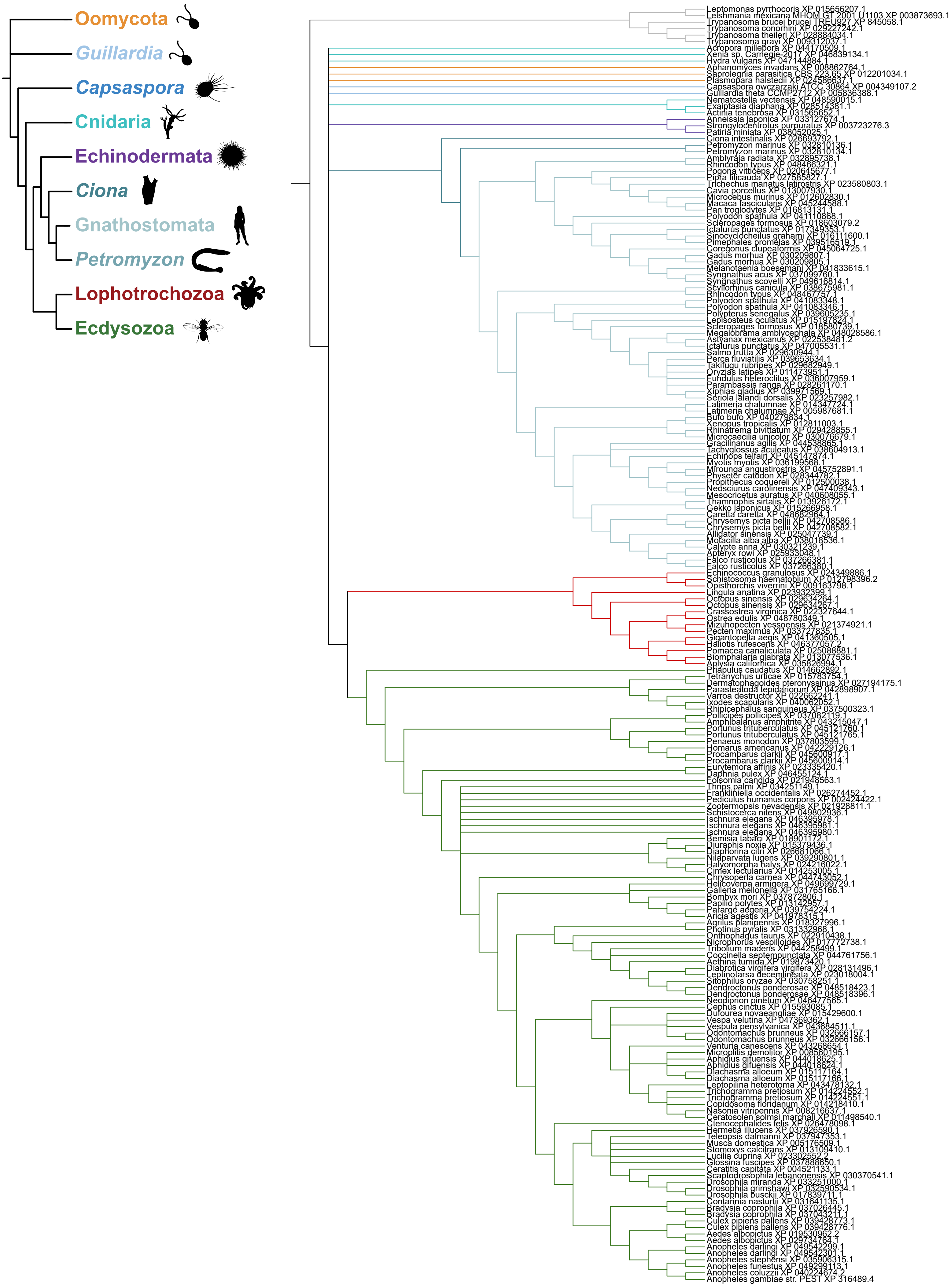

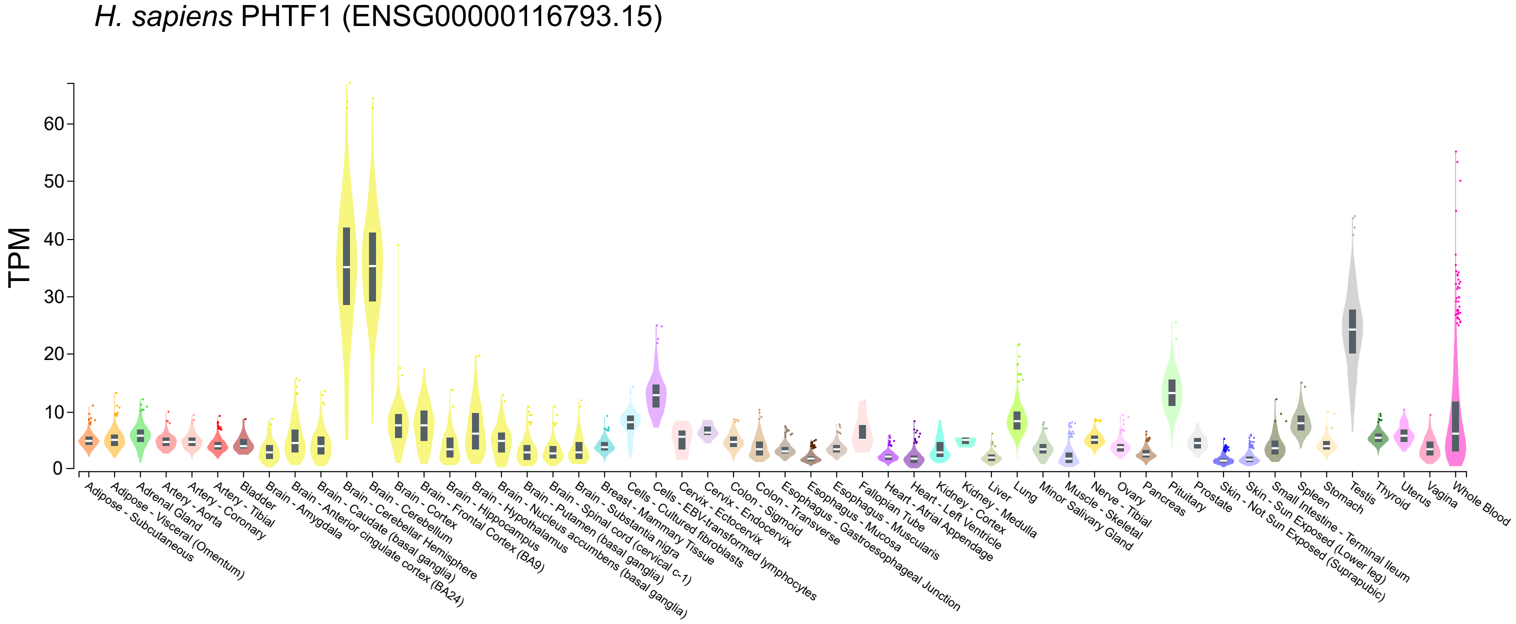
H. sapiens PHTF1 (ENSG00000116793.15)
TPM
0
10
20
30
40
50
60
Adipose - Subcutaneous
Adipose - Visceral (Omentum)
Adrenal Gland
Artery - Aorta
Artery - Coronary
Artery - Tibial
Bladder
Brain - Amygdala
Brain - Anterior cingulate cortex (BA24)
Brain - Caudate (basal ganglia)
Brain - Cerebellar Hemisphere
Brain - Cerebellum
Brain - Cortex
Brain - Frontal Cortex (BA9)
Brain - Hippocampus
Brain - Hypothalamus
Brain - Nucleus accumbens (basal ganglia)
Brain - Putamen (basal ganglia)
Brain - Spinal cord (cervical c-1)
Brain - Substantia nigra
Breast - Mammary Tissue
Cells - Cultured fibroblasts
Cells - EBV-transformed lymphocytes
Cervix - Ectocervix
Cervix - Endocervix
Colon - Sigmoid
Colon - Transverse
Esophagus - Gastroesophageal Junction
Esophagus - Mucosa
Esophagus - Muscularis
Fallopian Tube
Heart - Atrial Appendage
Heart - Left Ventricle
Kidney - Cortex
Kidney - Medulla
Liver
Lung
Minor Salivary Gland
Muscle - Skeletal
Nerve - Tibial
Ovary
Pancreas
Pituitary
Prostate
Skin - Not Sun Exposed (Suprapubic)
Skin - Sun Exposed (Lower leg)
Small Intestine - Terminal Ileum
Spleen
Stomach
Testis
Thyroid
Uterus
Vagina
Whole Blood

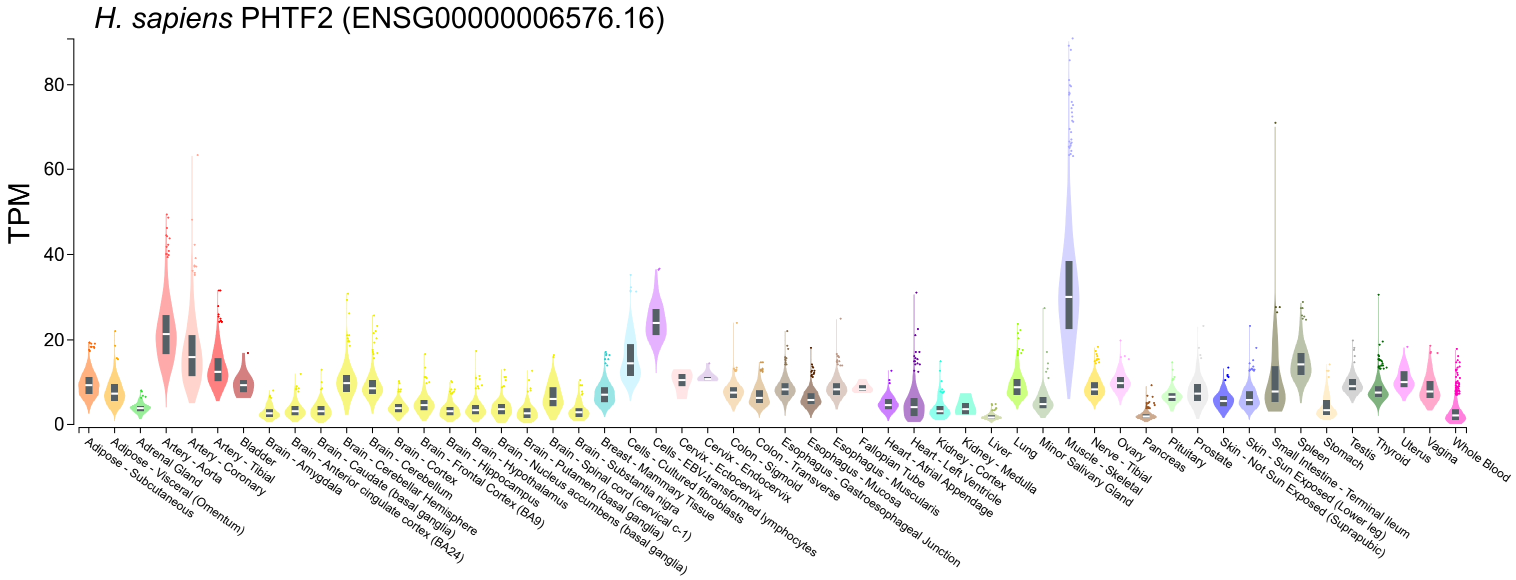
H. sapiens PHTF2 (ENSG00000006576.16)
TPM
0
20
40
60
80
Adipose - Subcutaneous
Adipose - Visceral (Omentum)
Adrenal Gland
Artery - Aorta
Artery - Coronary
Artery - Tibial
Bladder
Brain - Amygdala
Brain - Anterior cingulate cortex (BA24)
Brain - Caudate (basal ganglia)
Brain - Cerebellar Hemisphere
Brain - Cerebellum
Brain - Cortex
Brain - Frontal Cortex (BA9)
Brain - Hippocampus
Brain - Hypothalamus
Brain - Nucleus accumbens (basal ganglia)
Brain - Putamen (basal ganglia)
Brain - Spinal cord (cervical c-1)
Brain - Substantia nigra
Breast - Mammary Tissue
Cells - Cultured fibroblasts
Cells - EBV-transformed lymphocytes
Cervix - Ectocervix
Cervix - Endocervix
Colon - Sigmoid
Colon - Transverse
Esophagus - Gastroesophageal Junction
Esophagus - Mucosa
Esophagus - Muscularis
Fallopian Tube
Heart - Atrial Appendage
Heart - Left Ventricle
Kidney - Cortex
Kidney - Medulla
Liver
Lung
Minor Salivary Gland
Muscle - Skeletal
Nerve - Tibial
Ovary
Pancreas
Pituitary
Prostate
Skin - Not Sun Exposed (Suprapubic)
Skin - Sun Exposed (Lower leg)
Small Intestine - Terminal Ileum
Spleen
Stomach
Testis
Thyroid
Uterus
Vagina
Whole Blood

| | | *Phtf* | *Grl36a* | *Grl36b* | *Grl40a* | *Grl43a* | *Grl58a* | *Grl62a* | *Grl62b* | *Grl62c* | *Grl65a* | *GrlHz(A)* | *GrlHz(C)* |
|---|---|---|---|---|---|---|---|---|---|---|---|---|---|
| Adult male | Head | 8.19 | 0.06 | 2.54 | 0.03 | 0.06 | 0.09 | 0.07 | 0.03 | 0.00 | 0.90 | 3.20 | 0.22 |
| | Eye | 11.53 | 0.00 | 3.29 | 0.00 | 0.02 | 0.00 | 0.00 | 0.01 | 0.04 | 0.48 | 1.75 | 1.37 |
| | Brain | 15.33 | 0.09 | 5.14 | 0.00 | 0.02 | 0.17 | 0.00 | 0.03 | 0.01 | 0.05 | 1.99 | 0.82 |
| | Thoracicoabdominal ganglion | 10.32 | 0.04 | 5.30 | 0.03 | 0.01 | 0.28 | 0.01 | 0.02 | 0.02 | 0.18 | 3.39 | 0.96 |
| | Crop | 4.39 | 0.00 | 0.37 | 0.00 | 0.06 | 0.00 | 0.00 | 0.02 | 0.00 | 0.04 | 1.38 | 0.00 |
| | Midgut | 4.78 | 0.00 | 0.27 | 0.00 | 0.06 | 0.00 | 0.00 | 0.00 | 0.00 | 0.00 | 0.01 | 1.30 |
| | Hindgut | 5.2 | 0.00 | 0.16 | 0.00 | 0.01 | 0.01 | 0.01 | 0.00 | 0.00 | 0.00 | 0.36 | 0.39 |
| | Malpighian Tubules | 5.41 | 0.00 | 0.30 | 0.00 | 0.01 | 0.00 | 0.00 | 0.00 | 0.00 | 0.00 | 2.03 | 0.77 |
| | Fat body | 4.63 | 0.00 | 0.29 | 0.35 | 0.00 | 0.00 | 0.00 | 0.00 | 0.00 | 1.37 | 0.49 | 1.09 |
| | Salivary gland | 4.91 | 0.02 | 0.31 | 0.02 | 0.12 | 0.00 | 0.00 | 0.00 | 0.00 | 0.18 | 2.64 | 0.10 |
| | Heart | 10.29 | 0.00 | 0.23 | 0.00 | 0.04 | 0.00 | 0.00 | 0.00 | 0.00 | 0.64 | 8.76 | 0.00 |
| | Testis | 104.27 | 0.00 | 1.32 | 1.05 | 0.03 | 0.03 | 0.00 | 0.00 | 0.07 | 0.13 | 9.04 | 1.00 |
| | Accessory glands | 4.48 | 0.00 | 0.17 | 0.00 | 0.21 | 0.00 | 0.00 | 0.00 | 0.00 | 0.08 | 1.04 | 0.00 |
| | Carcass | 5.71 | 0.00 | 1.01 | 0.04 | 0.04 | 0.05 | 0.00 | 0.00 | 0.00 | 0.15 | 2.54 | 0.11 |
| | Rectal pad | 4.26 | 0.00 | 0.26 | 0.00 | 0.01 | 0.00 | 0.00 | 0.00 | 0.00 | 0.00 | 0.40 | 0.50 |
| | Whole body | 16.35 | 0.00 | 1.28 | 0.06 | 0.08 | 0.11 | 0.03 | 0.04 | 0.00 | 0.09 | 2.52 | 0.23 |
| Adult female | Head | 7.71 | 0.02 | 2.49 | 0.00 | 0.00 | 0.01 | 0.06 | 0.06 | 0.01 | 1.03 | 3.73 | 0.83 |
| | Eye | 9.02 | 0.00 | 2.38 | 0.00 | 0.00 | 0.00 | 0.00 | 0.00 | 0.01 | 0.50 | 3.98 | 0.67 |
| | Brain | 13.3 | 0.07 | 5.47 | 0.02 | 0.01 | 0.12 | 0.02 | 0.00 | 0.03 | 0.01 | 2.77 | 0.32 |
| | Thoracicoabdominal ganglion | 11.08 | 0.05 | 5.55 | 0.07 | 0.02 | 0.33 | 0.03 | 0.01 | 0.01 | 0.19 | 4.86 | 0.38 |
| | Crop | 3.34 | 0.00 | 0.16 | 0.00 | 0.01 | 0.02 | 0.00 | 0.00 | 0.00 | 0.02 | 1.93 | 0.00 |
| | Midgut | 2.75 | 0.00 | 0.09 | 0.00 | 0.00 | 0.00 | 0.00 | 0.00 | 0.00 | 0.00 | 0.17 | 0.60 |
| | Hindgut | 5.63 | 0.00 | 0.16 | 0.00 | 0.02 | 0.00 | 0.03 | 0.00 | 0.01 | 0.06 | 1.11 | 0.06 |
| | Malpighian Tubules | 4.74 | 0.00 | 0.42 | 0.01 | 0.01 | 0.00 | 0.00 | 0.02 | 0.00 | 0.02 | 1.35 | 0.31 |
| | Fat body | 3.69 | 0.00 | 0.08 | 0.00 | 0.07 | 0.00 | 0.00 | 0.00 | 0.00 | 3.25 | 4.53 | 0.00 |
| | Salivary gland | 4.54 | 0.00 | 0.12 | 0.02 | 0.17 | 0.02 | 0.00 | 0.00 | 0.02 | 0.08 | 5.94 | 0.16 |
| | Heart | 10.83 | 0.00 | 0.38 | 0.00 | 0.00 | 0.00 | 0.00 | 0.00 | 0.00 | 1.14 | 26.40 | 0.15 |
| | Ovary | 6.86 | 0.00 | 0.10 | 0.01 | 0.02 | 0.00 | 0.14 | 0.11 | 0.16 | 0.00 | 22.91 | 0.19 |
| | Virgin Spermatheca | 2.63 | 0.00 | 0.08 | 0.00 | 0.00 | 0.00 | 0.00 | 0.00 | 0.00 | 2.58 | 2.10 | 0.45 |
| | Mated Spermatheca | 2.86 | 0.00 | 0.00 | 0.00 | 0.05 | 0.00 | 0.00 | 0.00 | 0.00 | 3.12 | 2.96 | 0.00 |
| | Carcass | 5.57 | 0.00 | 0.68 | 0.00 | 0.01 | 0.00 | 0.00 | 0.00 | 0.00 | 0.15 | 6.20 | 0.58 |
| | Rectal pad | 4.43 | 0.00 | 0.18 | 0.00 | 0.06 | 0.00 | 0.00 | 0.00 | 0.00 | 0.00 | 1.35 | 0.35 |
| | Whole body | 6.04 | 0.00 | 0.15 | 0.03 | 0.00 | 0.00 | 0.06 | 0.12 | 0.05 | 0.10 | 14.30 | 0.30 |
| Larva | CNS | 5.18 | 0.05 | 1.40 | 0.00 | 0.07 | 0.03 | 0.08 | 0.21 | 0.14 | 0.02 | 4.97 | 0.15 |
| | Midgut | 5.49 | 0.00 | 0.25 | 0.00 | 0.00 | 0.00 | 0.00 | 0.00 | 0.00 | 0.00 | 0.85 | 0.09 |
| | Hindgut | 3.5 | 0.00 | 0.15 | 0.00 | 0.00 | 0.00 | 0.00 | 0.00 | 0.00 | 0.00 | 0.82 | 0.73 |
| | Malpighian Tubules | 2.88 | 0.00 | 0.02 | 0.00 | 0.02 | 0.00 | 0.00 | 0.00 | 0.00 | 0.00 | 5.31 | 0.34 |
| | Fat body | 5.64 | 0.00 | 0.00 | 0.00 | 0.05 | 0.07 | 0.13 | 0.07 | 0.00 | 0.07 | 8.60 | 3.22 |
| | Salivary gland | 3.5 | 0.00 | 0.02 | 0.00 | 0.09 | 0.00 | 0.00 | 0.00 | 0.03 | 0.03 | 1.33 | 0.55 |
| | Trachea | 3.12 | 0.00 | 2.71 | 0.00 | 0.01 | 0.00 | 0.00 | 0.00 | 0.00 | 0.00 | 1.64 | 0.24 |
| | Carcass | 2.57 | 0.08 | 0.32 | 0.00 | 0.00 | 0.00 | 0.00 | 0.00 | 0.03 | 0.00 | 2.16 | 0.34 |
| | Garland cells | 5.8 | 0.00 | 0.42 | 0.00 | 0.00 | 0.00 | 0.00 | 0.00 | 0.00 | 0.00 | 36.62 | 1.00 |
| | Whole body | 3.1 | 0.00 | 0.22 | 0.00 | 0.03 | 0.00 | 0.00 | 0.00 | 0.00 | 0.05 | 3.86 | 0.64 |

FKPM (Phtf scale): 120, 80, 40, 0

FKPM (Grl scale): 40, 35, 30, 25, 20, 15, 10, 5, 0

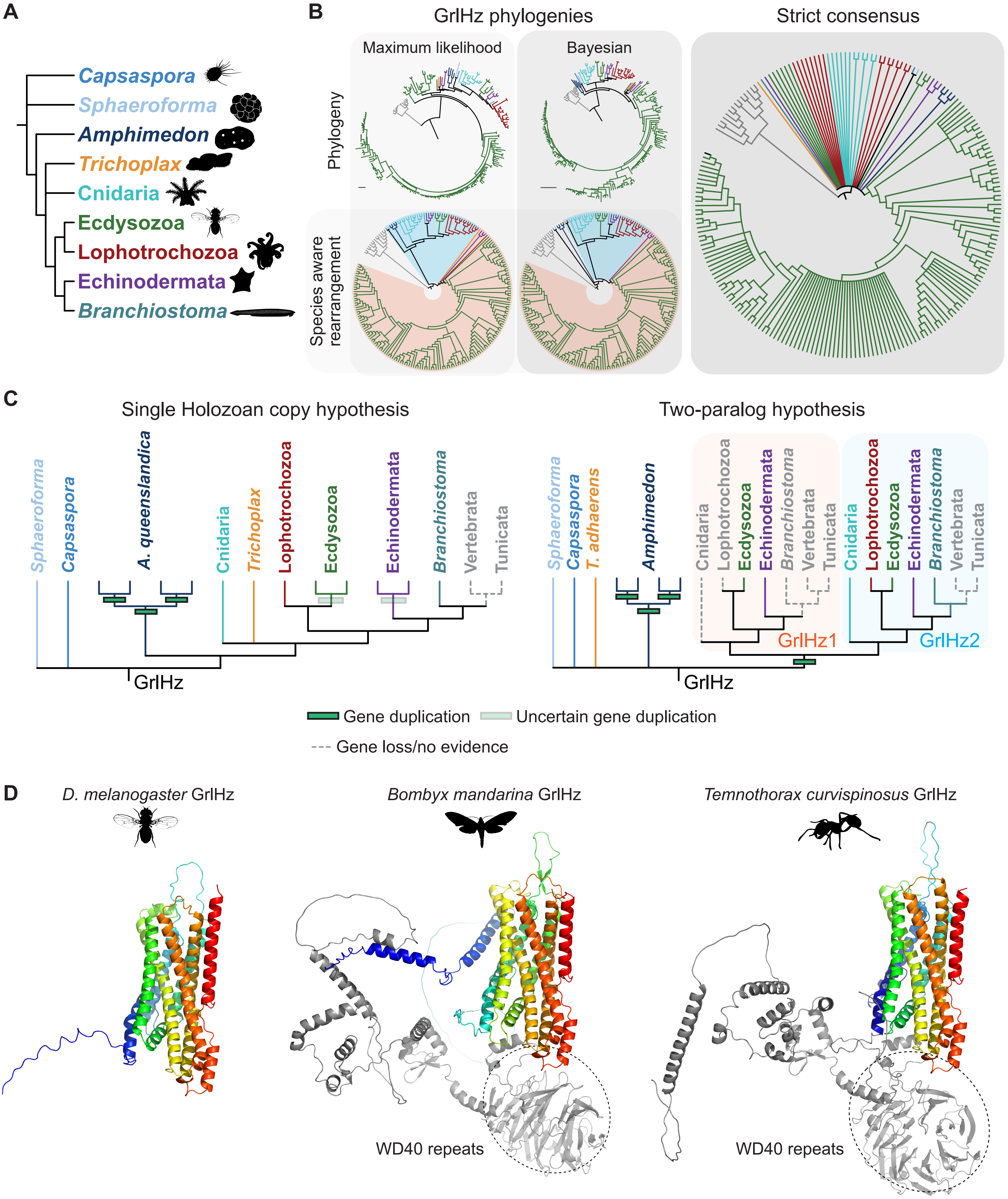
A
Capsaspora
Sphaeroforma
Amphimedon
Trichoplax
Cnidaria
Ecdysozoa
Lophotrochozoa
Echinodermata
Branchiostoma
B
GrlHz phylogenies
Maximum likelihood
Bayesian
Strict consensus
Phylogeny
Species aware rearrangement
C
Single Holozoan copy hypothesis
Sphaeroforma
Capsaspora
A. queenslandica
Cnidaria
Trichoplax
Lophotrochozoa
Ecdysozoa
Echinodermata
Branchiostoma
Vertebrata
Tunicata
GrlHz
Two-paralog hypothesis
Sphaeroforma
Capsaspora
T. adhaerens
Amphimedon
Cnidaria
Lophotrochozoa
Ecdysozoa
Echinodermata
Branchiostoma
Vertebrata
Tunicata
GrlHz1
Cnidaria
Lophotrochozoa
Ecdysozoa
Echinodermata
Branchiostoma
Vertebrata
Tunicata
GrlHz2
GrlHz
Gene duplication
Uncertain gene duplication
Gene loss/no evidence
D
D. melanogaster GrlHz
Bombyx mandarina GrlHz
Temnothorax curvispinosus GrlHz
WD40 repeats
WD40 repeats

Figure 3—figure supplement 2

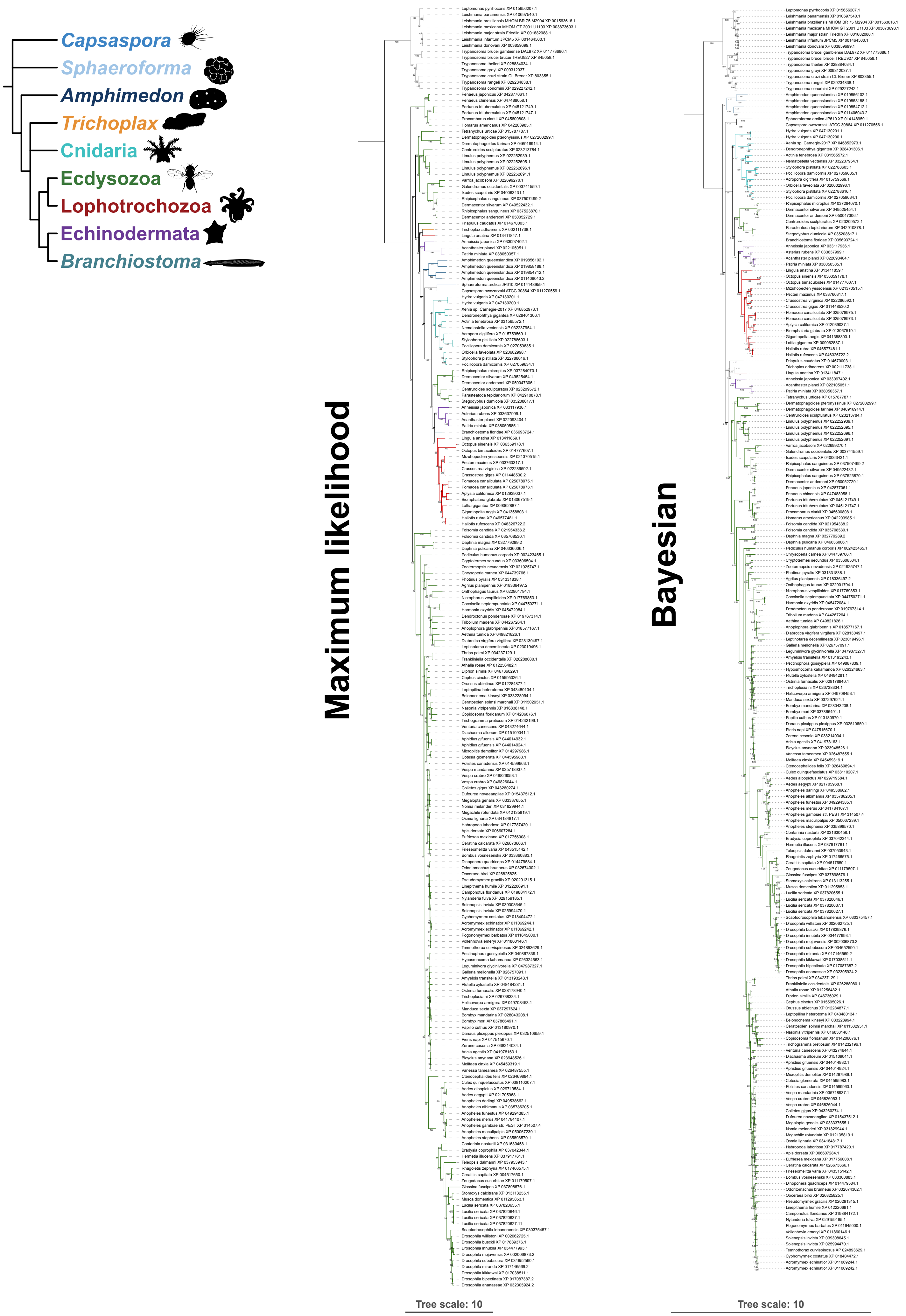

Figure 3—figure supplement 3

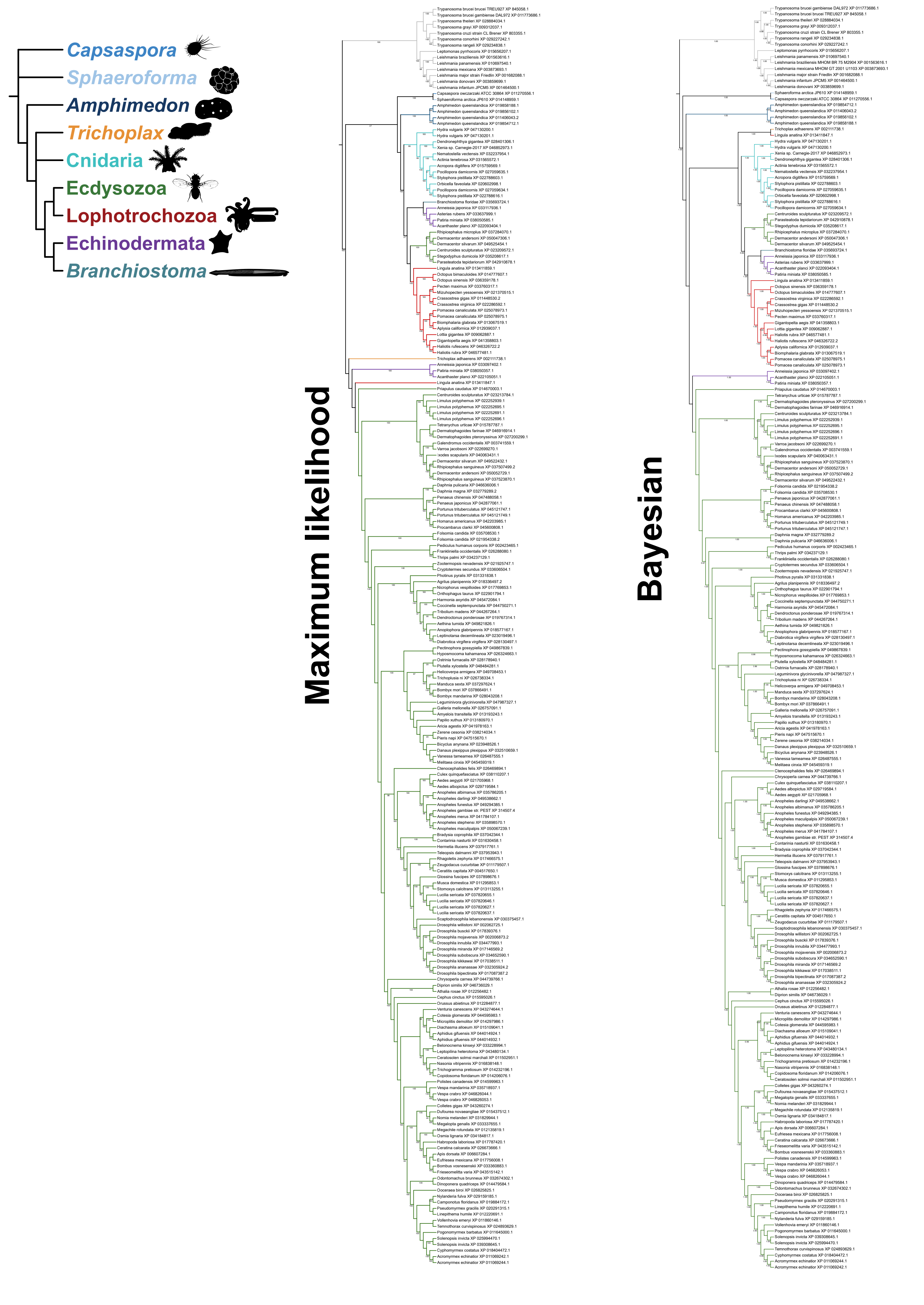

Figure 3—figure supplement 4

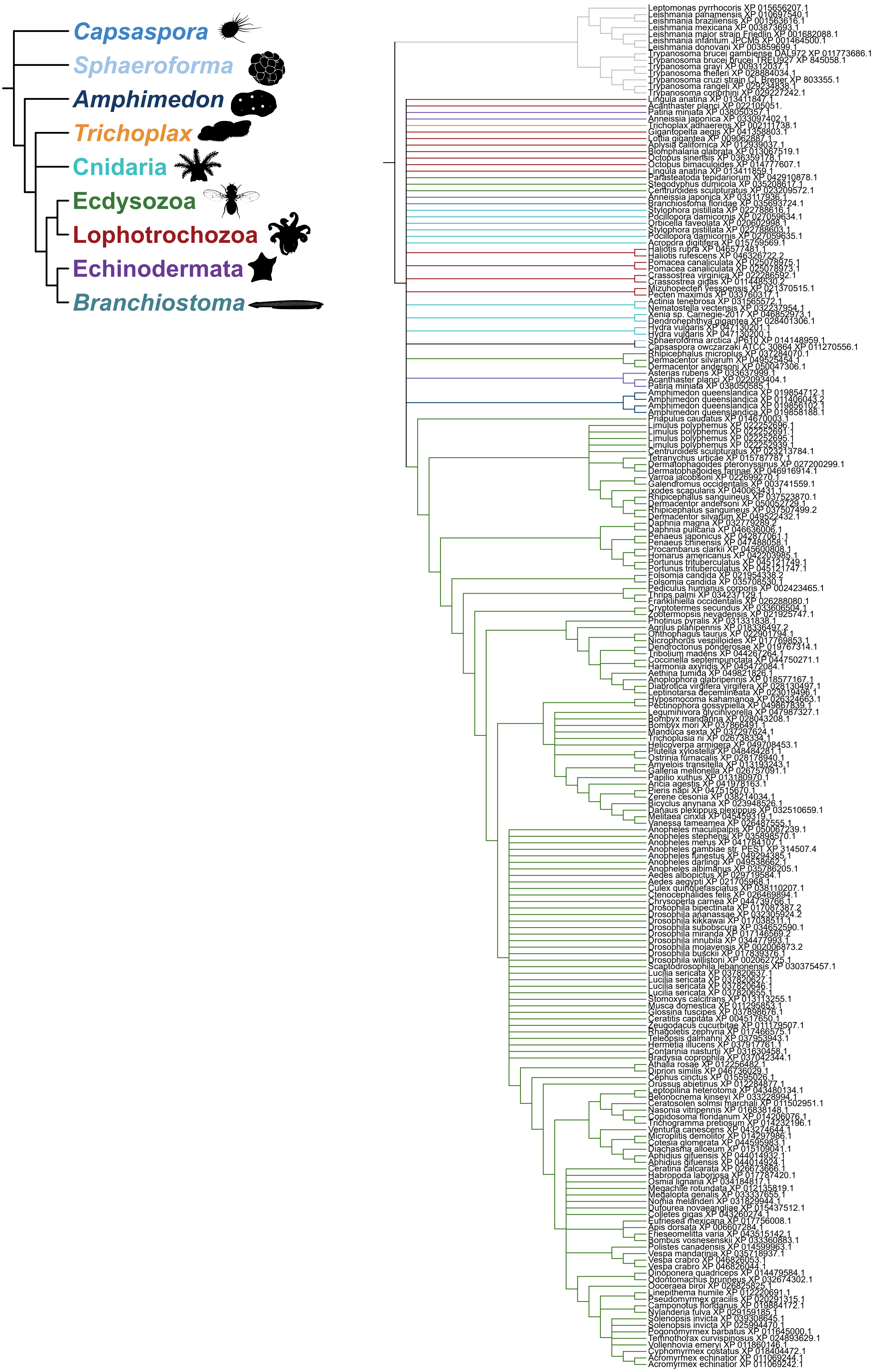

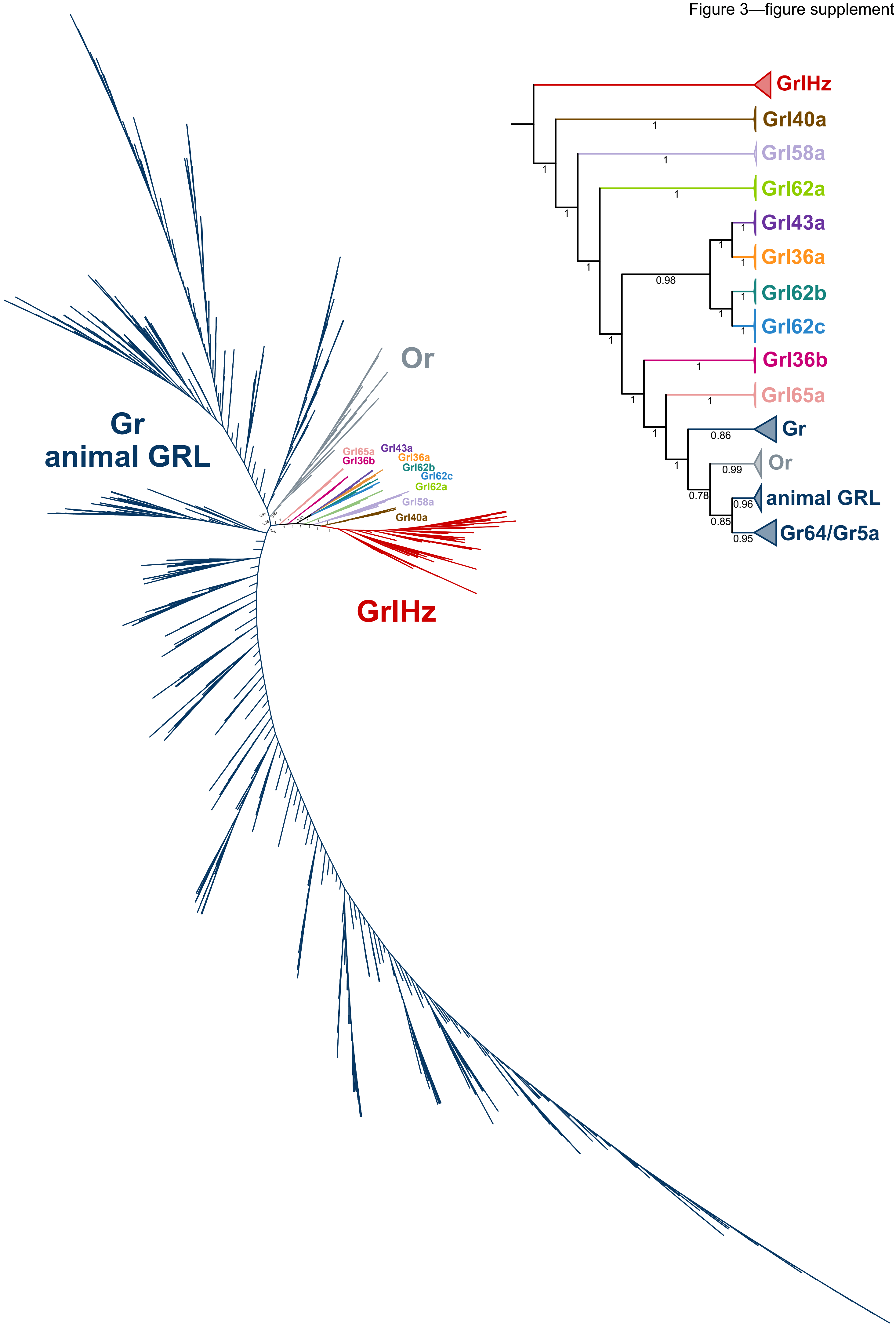
Or
Gr
animal GRL
GrlHz
Grl65a
Grl36b
Grl43a
Grl36a
Grl62b
Grl62c
Grl62a
Grl58a
Grl40a
GrlHz
Grl40a
Grl58a
Grl62a
Grl43a
Grl36a
Grl62b
Grl62c
Grl36b
Grl65a
Gr
Or
animal GRL
Gr64/Gr5a
1
0.98
0.86
0.99
0.78
0.96
0.85
0.95

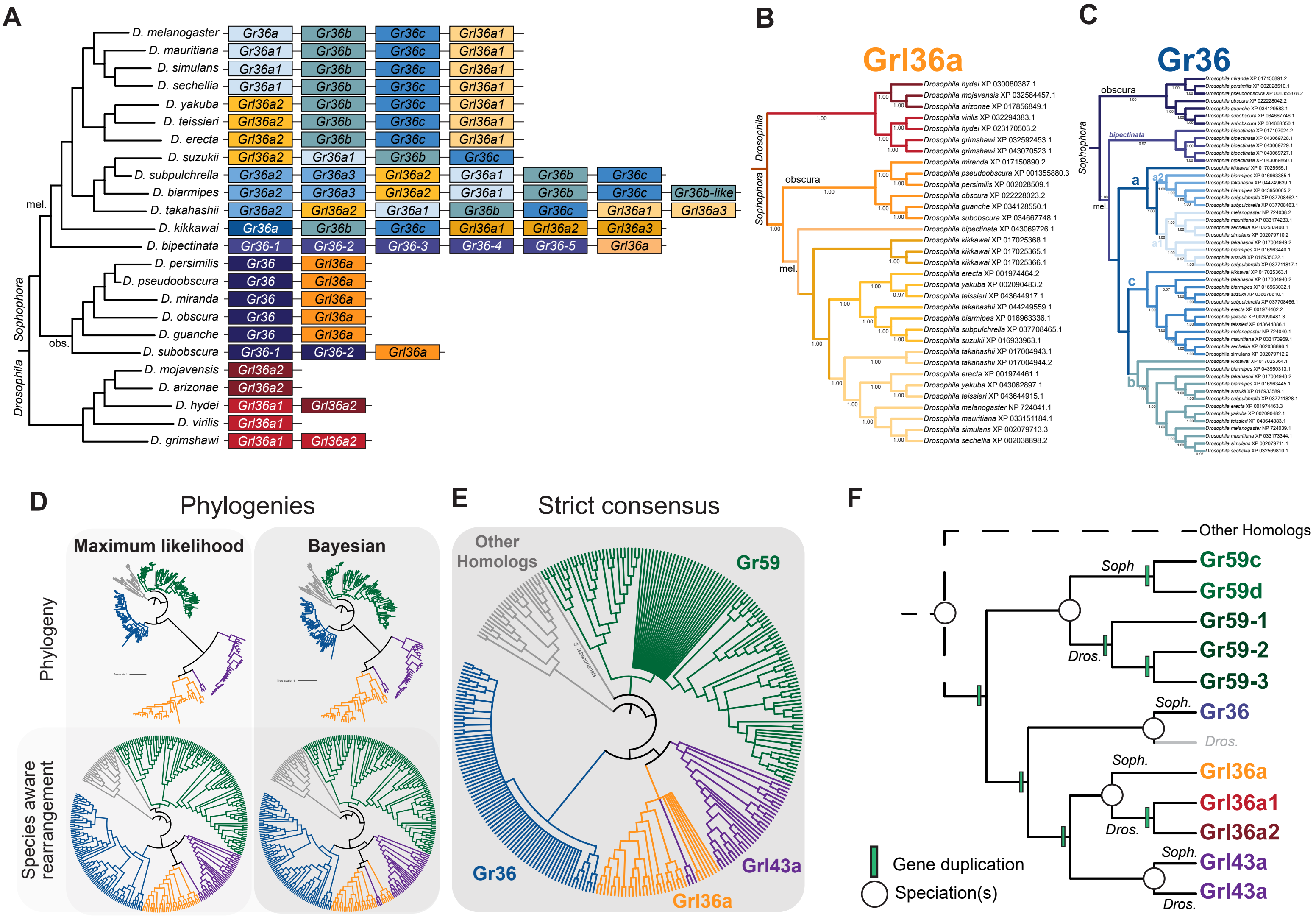
A
D. melanogaster
D. mauritiana
D. simulans
D. sechellia
D. yakuba
D. teissieri
D. erecta
D. suzukii
D. subpulchrella
D. biarmipes
D. takahashii
D. kikkawai
D. bipectinata
D. persimilis
D. pseudoobscura
D. miranda
D. obscura
D. guanche
D. subobscura
D. mojavensis
D. arizonae
D. hydei
D. virilis
D. grimshawi
mel.
obs.
Sophophora
Drosophila
Gr36a
Gr36b
Gr36c
Grl36a1
Grl36a2
Grl36a3
Gr36a1
Gr36a2
Gr36a3
Gr36b-like
Gr36-1
Gr36-2
Gr36-3
Gr36-4
Gr36-5
Grl36a
Gr36
B
Grl36a
Drosophila
Sophophora
obscura
mel.
C
Gr36
obscura
bipectinata
Sophophora
mel.
a
a2
a1
c
b
D
Phylogenies
Maximum likelihood
Bayesian
Phylogeny
Species aware rearrangement
E
Strict consensus
Other Homologs
Gr59
Gr36
Grl36a
Grl43a
F
Other Homologs
Soph
Gr59c
Gr59d
Dros.
Gr59-1
Gr59-2
Gr59-3
Soph.
Gr36
Dros.
Soph.
Grl36a
Grl36a1
Dros.
Grl36a2
Soph.
Grl43a
Grl43a
Dros.
Gene duplication
Speciation(s)

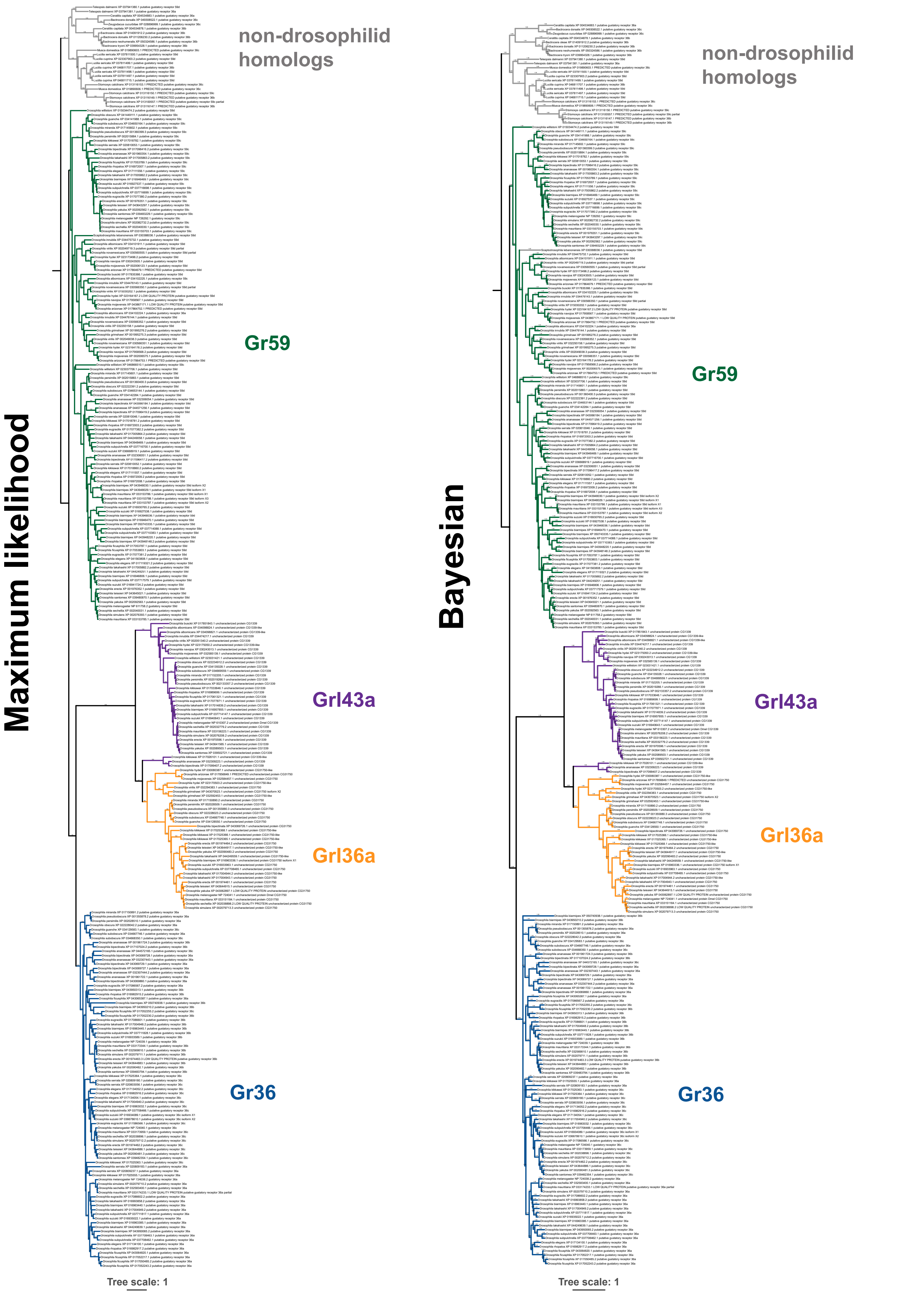
Maximum likelihood
non-drosophilid homologs
Gr59
Grl43a
Grl36a
Gr36
Tree scale: 1
Bayesian
non-drosophilid homologs
Gr59
Grl43a
Grl36a
Gr36
Tree scale: 1

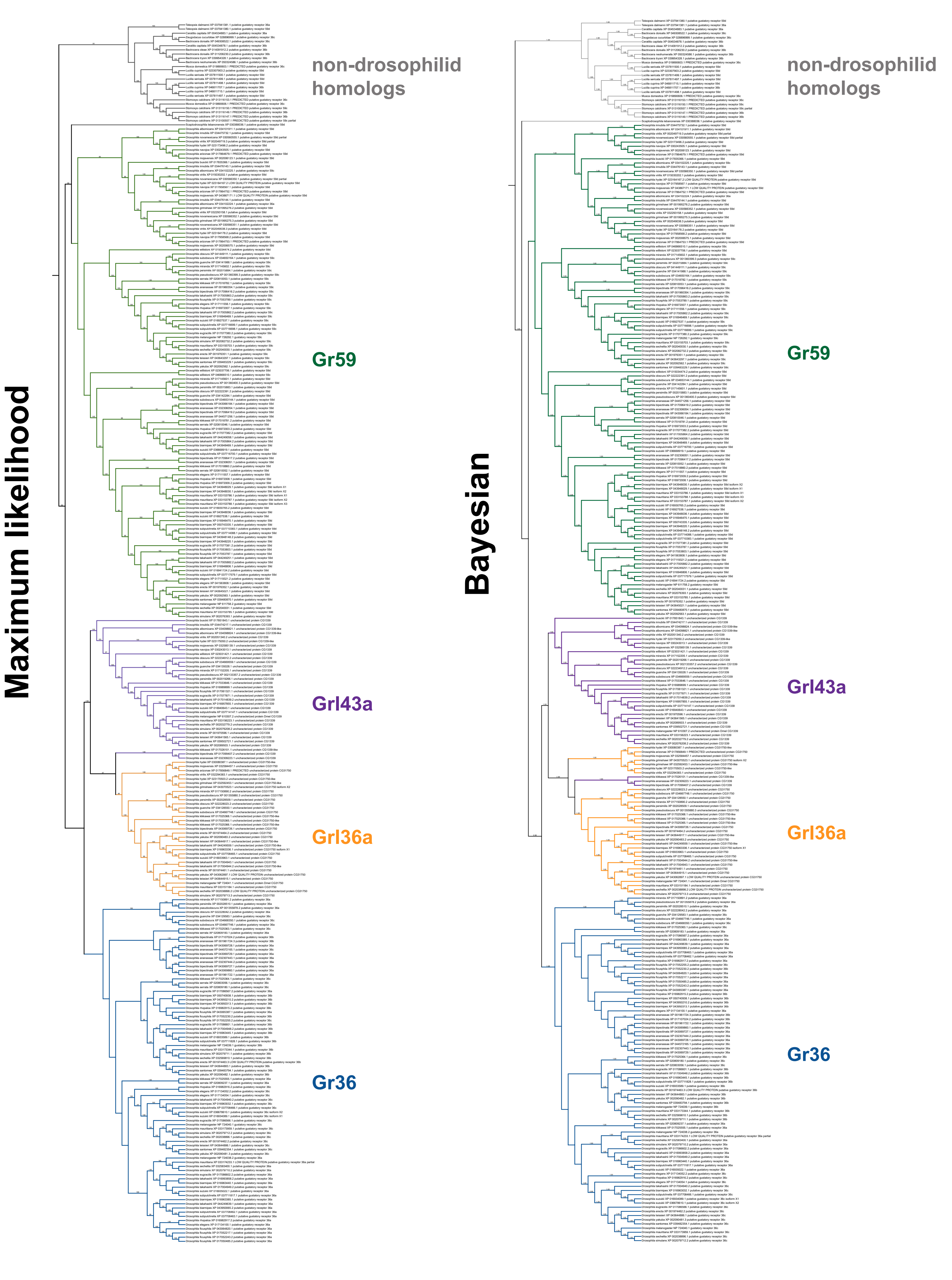
Maximum likelihood
non-drosophilid homologs
Gr59
Grl43a
Grl36a
Gr36
Bayesian
non-drosophilid homologs
Gr59
Grl43a
Grl36a
Gr36

Figure 3—figure supplement 9

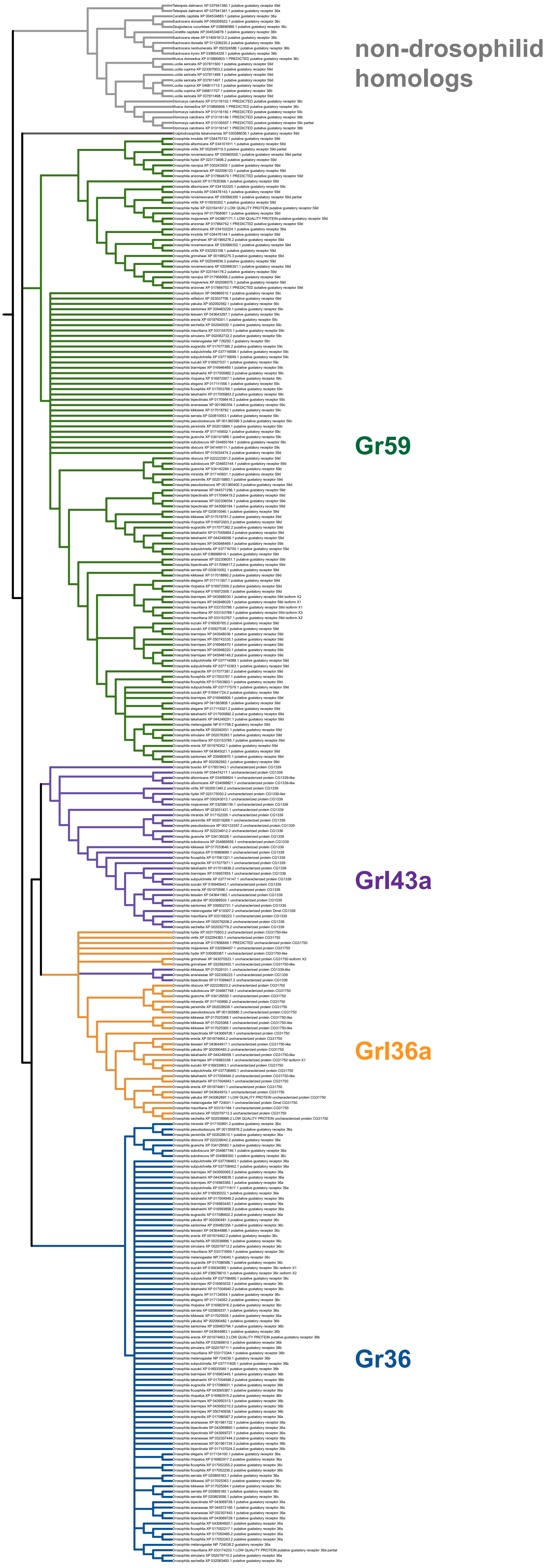

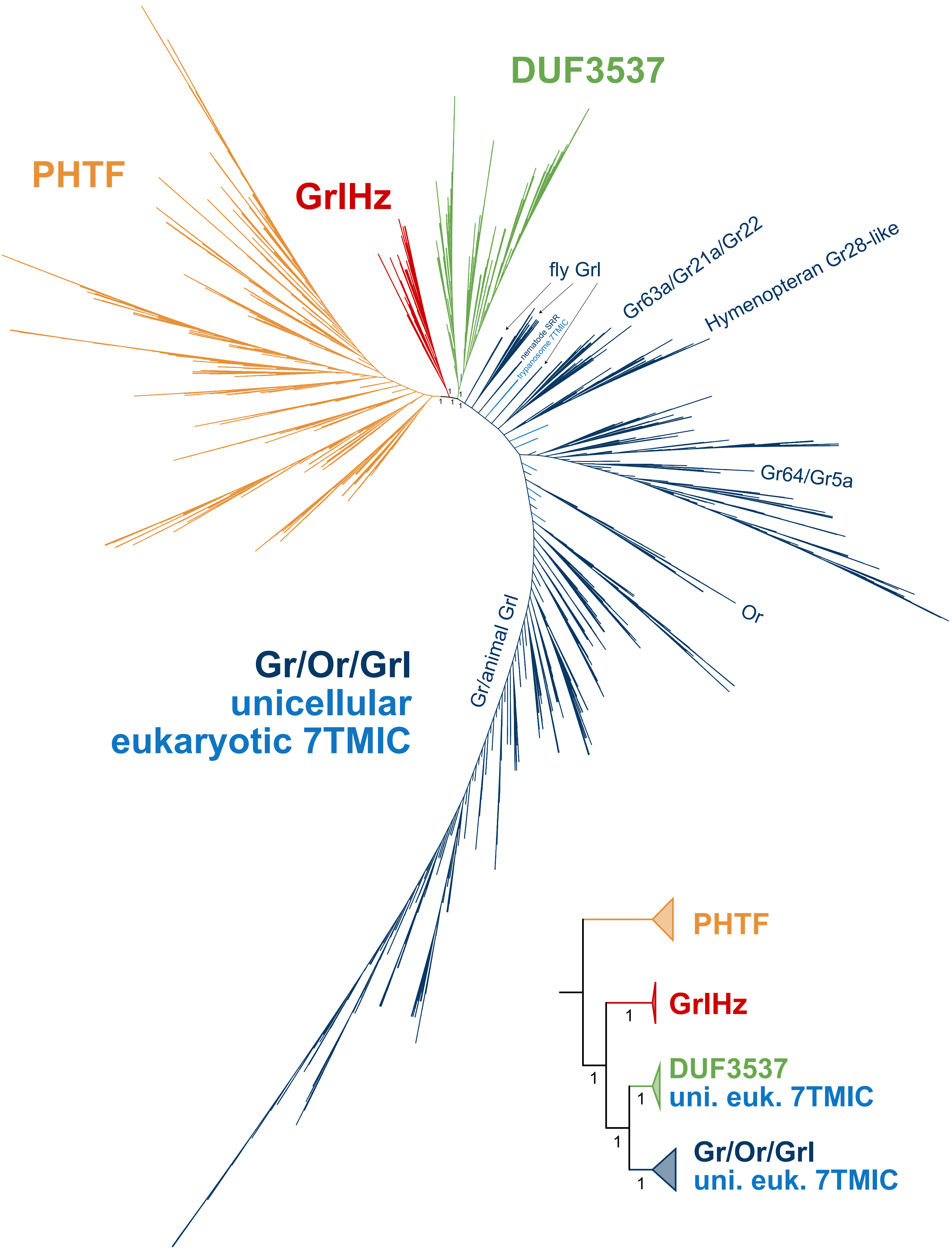
PHTF
GrlHz
DUF3537
fly Grl
nematode SRR
trypanosome 7TMIC
Gr63a/Gr21a/Gr22
Hymenopteran Gr28-like
Gr64/Gr5a
Or
Gr/animal Grl
Gr/Or/Grl
unicellular
eukaryotic 7TMIC
PHTF
GrlHz
DUF3537
uni. euk. 7TMIC
Gr/Or/Grl
uni. euk. 7TMIC
